# Super-relaxed state of myosin in human skeletal muscle is fiber-type dependent

**DOI:** 10.1101/2020.09.24.311795

**Authors:** Lien A. Phung, Aurora D. Foster, Mark S. Miller, Dawn A. Lowe, David D. Thomas

## Abstract

The myosin super-relaxed state (SRX) in skeletal muscle is hypothesized to play an important role in regulating muscle contractility and thermogenesis in humans, but has only been examined in model organisms. Here we report the first human skeletal muscle SRX measurements, using quantitative epifluorescence microscopy of fluorescent 2’/3’-O-(N-methylanthraniloyl) ATP (mantATP) single-nucleotide turnover. Myosin heavy chain (MHC) isoform expression was determined using gel electrophoresis for each permeabilized vastus lateralis fiber, to allow for novel comparisons of SRX between fiber-types. We find that the fraction of myosin in SRX is less in MHC IIA fibers than in MHC I and IIAX fibers (p = 0.008). ATP turnover of SRX is faster in MHC IIAX fibers compared to MHC I and IIA fibers (p = 0.001). We conclude that SRX biochemistry is measurable in human skeletal muscle, and our data indicate that SRX depends on fiber type as classified by MHC isoform. Extension from this preliminary work would provide further understanding regarding the role of SRX in human muscle physiology.

## Introduction

Skeletal muscle has two principal functional states: relaxation and active force production. Decades of molecular research focused on the active state have dissected the myosin-actin interactions that drive force production. Advances in biophysical tools in the past decade have inspired further investigations into the temporally more prevalent relaxed state, in which myosin is detached from actin and exists in two distinct substates: disordered relaxed (DRX) and super-relaxed (SRX) (13). SRX is characterized by extremely slow ATP turnover kinetics due to autoinhibition of the myosin catalytic domains (13). The existence of DRX and SRX is conserved among model organisms including rabbits, mice, and tarantulas (4, 17, 23, 27). The ubiquity of SRX suggests that this substate has physiological importance in human health, such as modulation of resting muscle metabolic rate (19) and contribution to sex-specific, age-related muscle weakness (23). Therefore, investigating these relaxation substates in human skeletal muscle is important, as myosin isoforms behave differently based upon species (21).

SRX has not previously been measured in human muscle, nor has SRX been compared among fibers that express different myosin heavy chain (MHC) isoforms (otherwise known as fiber types). Adult human skeletal muscle fibers express MHC I, IIA, IIX isoforms (16). Some fibers co-express more than one MHC isoform to give rise to hybrid fiber types, such as IIAX (16). Phenotypically, fibers expressing different MHC isoforms can be classified as slow-fast via properties of contractile velocity (3), oxidative capacity, or fatigue resistance (22). MHC I fibers are considered slow-twitch fibers, whereas MHC II fibers are classically considered fast-twitch. Compared to MHC II fibers and associated subtypes, MHC I fibers have slower contractile velocity, lower power output, and less calcium sensitivity (3).

Muscles are composed of fibers containing one or more MHC isoforms, enabling substantial differences in contractile performance (21) due to their varied biochemical properties, such as active ATP turnover kinetics (9), which may extend to relaxation. There are >4-fold differences in ATPase activity among fibers containing different MHC isoforms during contraction (9, 21). If those large differences are limited to myosin ATPase function and not mechanical coupling properties, then ATP turnover rate differences are expected among fibers of various types during relaxation as well. Previous SRX work in rabbit muscle focused mainly on the fast-contracting psoas fibers (13, 27), which are predominantly MHC IIX (8), so the need to consider variable MHC expression was low. However, data suggests that slowly-contracting soleus fibers have different SRX biochemistry in terms of ATP turnover lifetime (27). Discerning the SRX biochemistry of different MHC isoforms is necessary to gain a better understanding of skeletal muscle function and metabolism. Therefore, the present study focuses on establishing the feasibility of measuring the SRX in human fibers and examining the differences among MHC isoforms. The objective of this study is to measure DRX and SRX in human skeletal muscle fibers and to distinguish characteristics among various fiber types identified across individuals independent of sex and age. Among MHC I/IIA/IIAX isoforms, we found that the fraction of SRX in MHC IIA fibers is relatively low and that nucleotide turnover lifetime is fast in IIAX fibers compared to the other two fiber types.

## Materials and methods

One young and three older adults provided biopsies for this study (**Fig. 1*A***). These volunteers were classified as sedentary (less than two 30-minute exercise sessions per week) by self-report. Written informed consent was obtained from each healthy (no history of neuromuscular disease) volunteer prior to local lidocaine anesthesia. Needle biopsies were obtained from the lateral portion of the vastus lateralis via incision 10 cm proximal to the knee and at a relatively constant depth within the muscle body as determined by the location of the muscle fascia. All procedures were approved by the University of Massachusetts Institutional Review Board (IRB #218 and #1014). Biopsy samples were dissected into bundles (4°C) and prepared for storage (−20°C) in 50% glycerol solution (28).

**Fig. 1.**
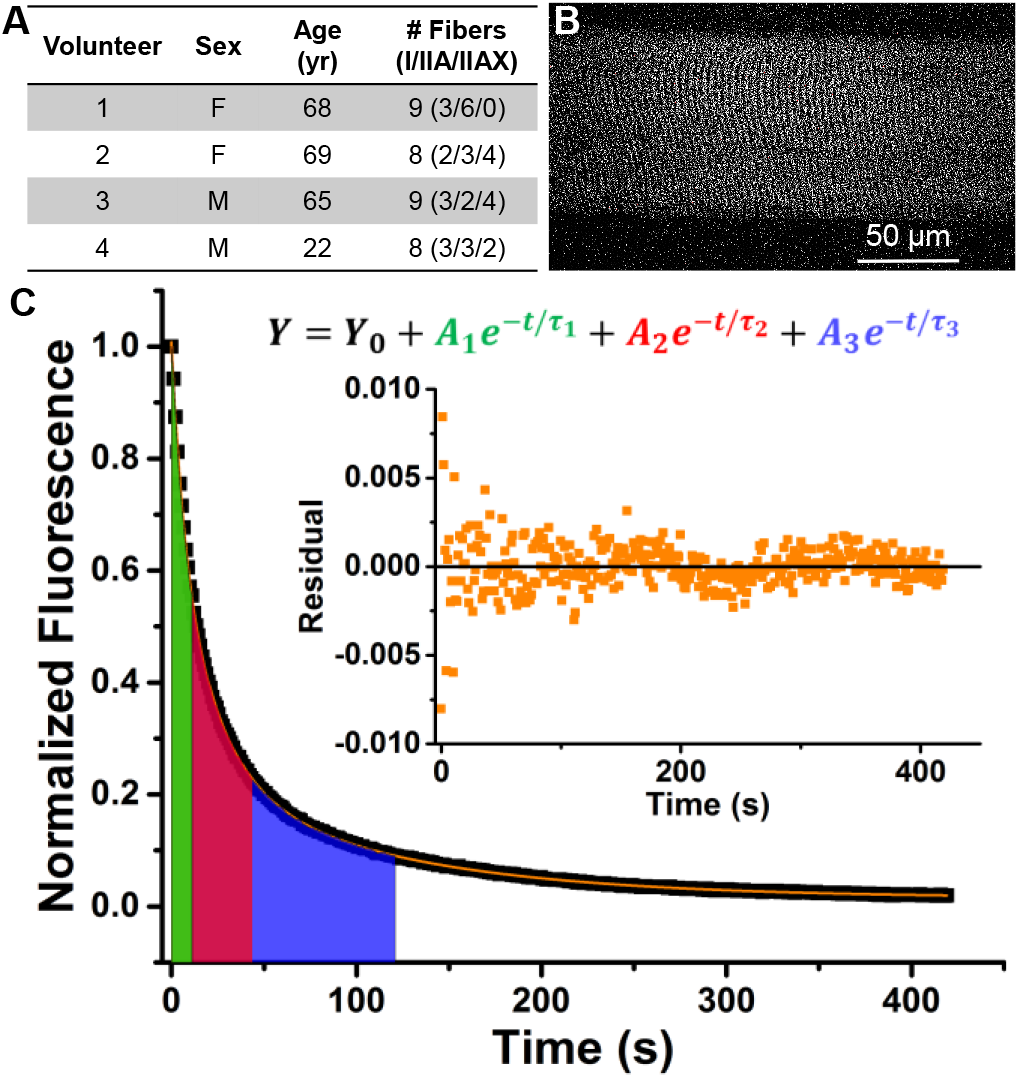
Measuring fluorescent nucleotide exchange activity of human skeletal muscle fibers. (A) Volunteer characteristics. (B) Representative fluorescence image of a single human skeletal muscle fiber loaded with mantATP prior to start of ATP exchange wash. (C) Representative normalized fluorescence trace (black) fitted with a 3-exponential decay function (orange). Color-shaded areas under the curve reflect the lifetime (τ) of each exponential decay component. Green: nonspecific washout; red: DRX; blue: SRX. Inset: Residual plot from fitting function.

Nucleotide exchange experiments were performed within 2 months of sample collection. On each experiment day, a muscle bundle was permeabilized with 1% Triton X-100 for 30 min (4°C). Single fibers were manually isolated and fluorescent nucleotide turnover activity was measured at 23 ± 1 °C and analyzed as previously described (23). Briefly, we used quantitative epifluorescence microscopy to analyze relaxation substates of myosin in those fibers. Time-lapsed images of fibers saturated with 250 μM mantATP and their fluorescence intensity were recorded after 4 mM non-fluorescent ATP was added to start nucleotide exchange (**Fig. 1*B***). During the experiment, fluorescence intensity decreased as bound mant-nucleotides were released from myosin and replaced by non-fluorescent ATP **(Fig. 1*C***). This process is effectively modeled by a 3-exponential decay function (23), where the fractions and lifetimes of DRX and SRX correspond to pre-exponential amplitudes (A) and exponential decay constants (τ) of the second (DRX) and third decay (SRX) components, respectively. A shorter lifetime represents a faster nucleotide turnover process of the respective myosin state. MHC isoform composition of each fiber was determined by SDS-PAGE (15). The number of fibers analyzed per type per subject is indicated in **Fig. 1*A***.

Average data are presented as mean ± SEM. The Shapiro-Wilk test was used to investigate within group normality for a given parameter of interest. Levene’s test was conducted to assess the homogeneity of the variance assumption. One-way analysis of variance and analysis of covariance were used to analyze fraction and lifetime data. Correlations were determined with Pearson’s correlation coefficient. Statistical analyses were carried out using OriginLab 2015 and IBM SPSS software. P-values ≤ 0.05 were considered statistically significant.

## Results

First we demonstrated, by epifluorescence microscopy, that the myosin SRX state in human skeletal muscle fibers is measurable in MHC I, IIA and IIAX fibers, the most common fiber-types in humans (16). The fraction of DRX was 0.57 ± 0.03 and 0.57 ± 0.05 in MHC I and IIAX fibers, which differed from MHC IIA fibers at 0.70 ± 0.03 (**Fig. 2*A***). Likewise, the fraction of SRX was 0.43 ± 0.03 and 0.43 ± 0.05 in MHC I and IIAX, which differed from MHC IIA at 0.30 ± 0.03 (**Fig. 2*B***). We conclude that the fraction of muscle myosin in each relaxed substate is dependent on the MHC isoform.

**Fig. 2.**
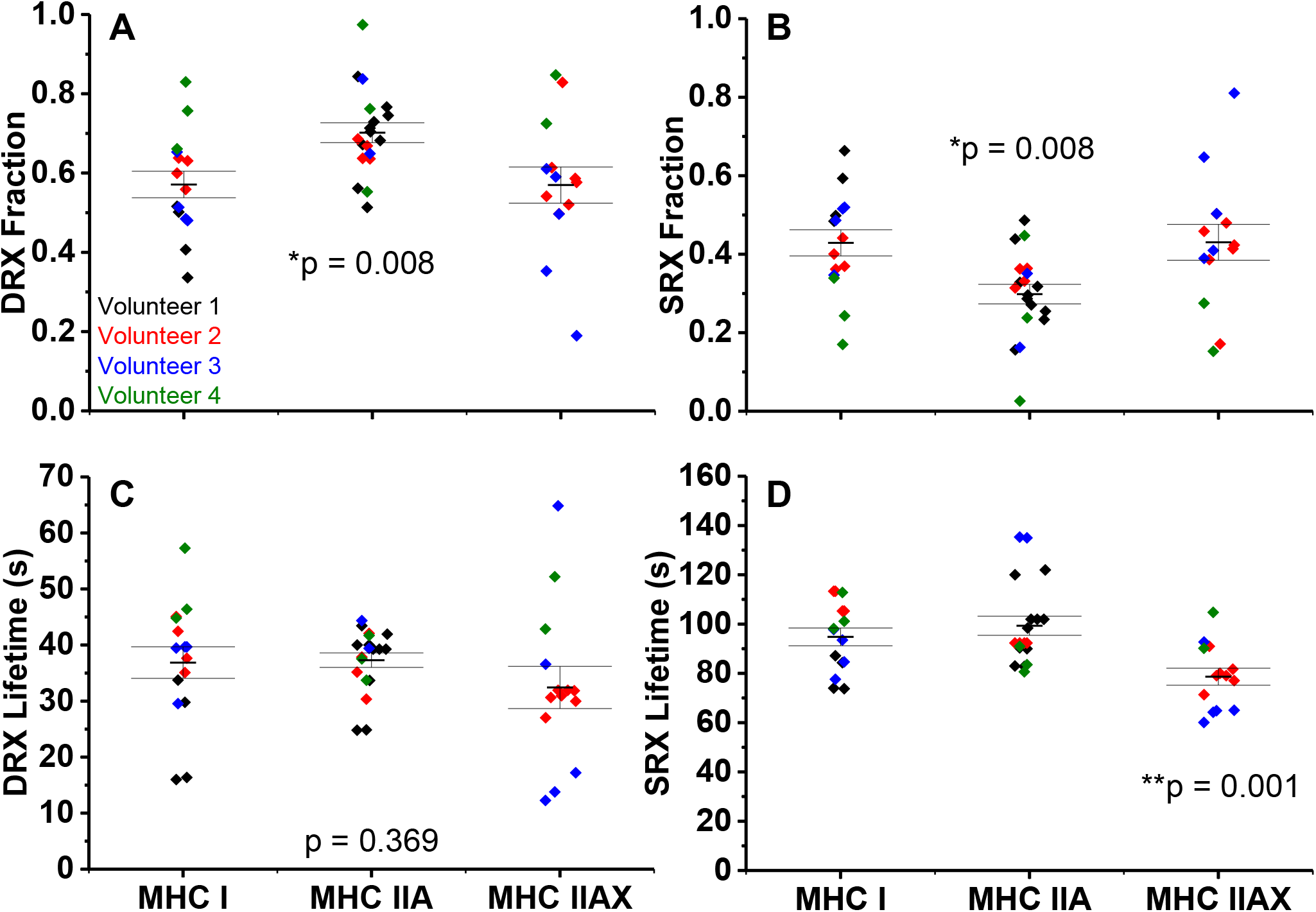
Relaxation states of skeletal muscle myosin in fibers isolated from human muscle. Fraction of myosin in DRX (A) and SRX (B). Lifetime (τ) of nucleotide turnover by myosin in DRX (C) and in SRX (D). Horizontal bars represent mean ± SEM. Individual symbols are values from each nucleotide exchange experiment for that fiber and color coded by volunteer. *MHC IIA differs from MHC I and IIAX, **MHC IIAX differs from MHC I and IIA as determined by one-way ANOVA.

The lifetime of nucleotide turnover, or ATP turnover kinetics, in DRX was not different between fiber-types, 37 ± 3, 37 ± 1 and 32 ± 4 s for MHC I, IIA and IIAX fibers (**Fig. 2*C***). Hence, the single ATP turnover kinetics of human skeletal muscle fibers in DRX is not fiber-type dependent. The lifetime of nucleotide turnover in SRX was 79 ± 3 s in MHC IIAX fibers, faster than the 95 ± 4 and 99 ± 4 s for MHC I and IIA fibers (**Fig. 2*D***). The ATP turnover kinetics of myosin SRX is significantly faster in MHC IIAX fibers compared to the other two MHC isoforms. Notably, MHC IIAX fibers contained 40-55% IIX isoform expression.

To determine if relaxation states were influenced by fiber morphology, we examined the relationships of DRX and SRX measurements with cross-sectional area (range: 1,571 to 11,165 μm^2^) or resting sarcomere length (range: 1.70 to 2.69 μm). There was no significant correlation between cross-sectional area and relaxation states fractions or lifetimes (p ≥ 0.15), suggesting that skeletal myosin relaxation substates are not dependent on fiber size. The fraction and lifetime of myosin DRX and SRX correlated with resting sarcomere length (p ≤ 0.006, (**Fig. 3*A-D***). The fraction of DRX is negatively associated with sarcomere length (**Fig. 3*A***), while the fraction of SRX is positively correlated with sarcomere length (**Fig. 3*B***). This indicates that in a resting fiber with longer sarcomere lengths, the conditions are more favorable for myosin heads to fold back and form the interacting heads motif (IHM) (20), the characteristic structural motif of SRX (13). The lifetimes of DRX and SRX were negatively associated with sarcomere length (**Fig. 3*C-D***), showing that the length of the time spent in DRX and SRX shortens as sarcomere length increases.

**Fig. 3.**
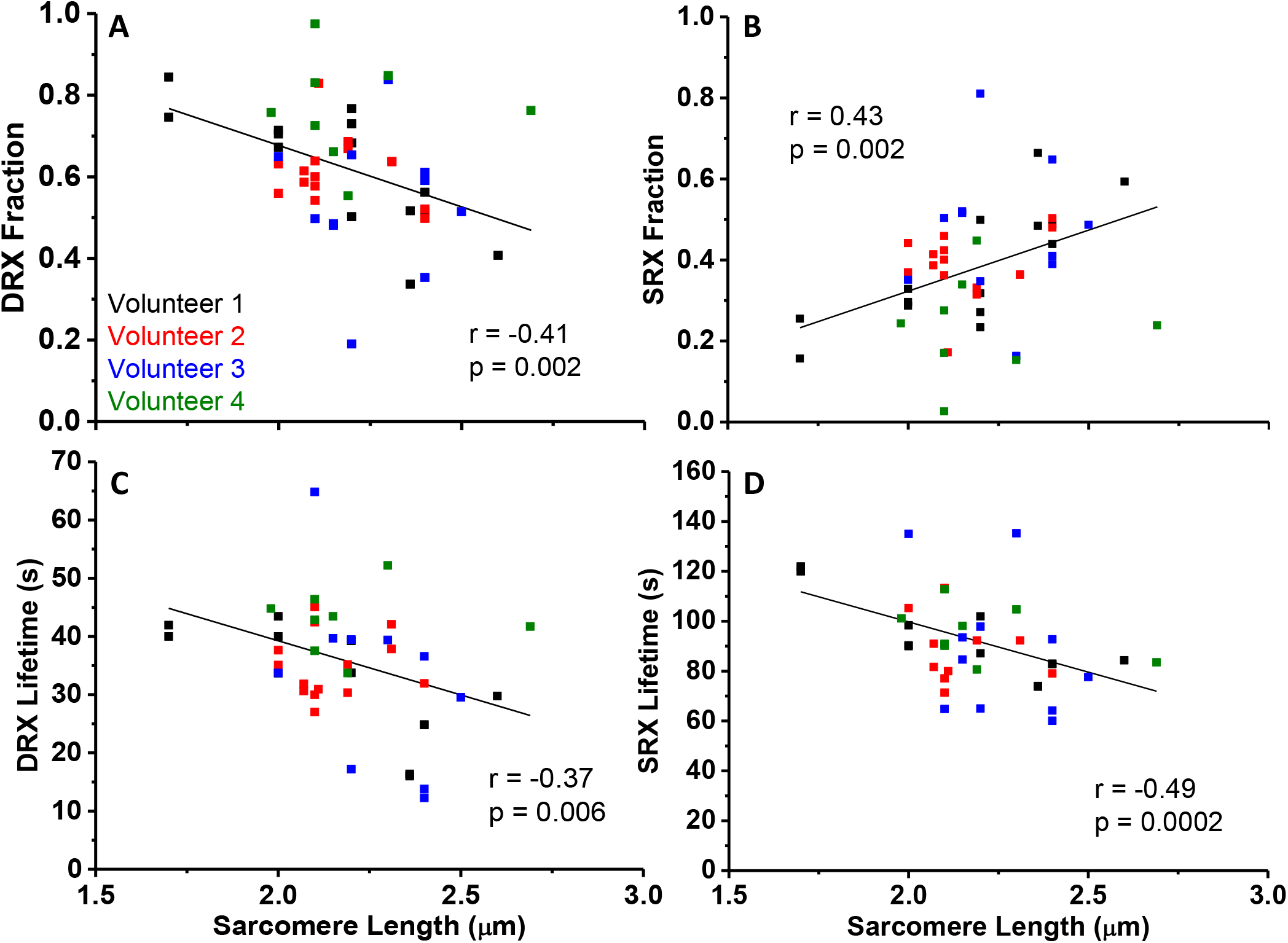
Relationship of fiber relax parameters and resting sarcomere length. Fraction of myosin in DRX (A) and SRX (B) in relationship to fiber sarcomere length. Lifetime (τ) of nucleotide turnover by myosin in DRX (C) and in SRX (D) related to fiber sarcomere length. Pearson correlation coefficient (r). Individual symbols are values from each nucleotide exchange experiment for that fiber and color coded by volunteer.

When adjusted for sarcomere length as a covariate, the conclusions regarding fiber type differences were unchanged. Fibers composed of MHC IIA had a lower fraction of SRX (higher fraction of DRX) compared to MHC I and IIAX fibers (p = 0.012). While DRX lifetime did not differ among fiber types (p = 0.48), SRX lifetime remained shorter in MHC IIAX than I and IIA fibers (p = 0.001).

## Discussion

This work demonstrates for the first time that human skeletal muscle fibers are characterized by distinct DRX and SRX substates, as previously observed for other vertebrate organisms (4, 13, 23). We also present evidence that skeletal muscle SRX properties are MHC isoform-dependent; which is likely to influence whole muscle metabolic rate and performance. From the weighted average ATPase rates of DRX and SRX of each fiber type, we calculated that MHC IIA and IIAX fibers consume ATP 10% and 17% faster than MHC I fibers, respectively. This higher resting energetic consumption of MHC II fibers may contribute to differences in whole-body basal metabolic rate - given that a 20% transition away from SRX would double muscle thermogenesis as reported by Stewart *et al*. (27). These new results provide additional important considerations for understanding muscle physiology, and enhance motivation for future in-depth studies of human skeletal muscle resting properties in all MHC isoforms.

While the differences in active ATPase rates among MHC isoforms are large (9, 21), rate differences in relaxation are small (**Fig. 2*C*&*D***). Active myosin ATPase is influenced by its thick filament arrangement and the resulting interaction with the thin filament. For example, thick filaments in MHC I and IIA fibers have been shown to have simple lattice spacing while MHC IIX/IIB/IID fibers have superlattice spacing in rats (12). The differential arrangement of thick filaments may impact the contractile properties of a fiber by affecting the efficiency of myosin head attachment to the actin thin filament in cross-bridge cycling, as previously proposed (12). Furthermore, the filament lattice arrangement perspective offers insights into how the duration of the active state in isometric twitch did not correlate with the ATPase activity of purified myosin (2). The lattice spacing arrangement may also affect the probability of a myosin head forming the auto-inhibited state effectively, which could explain the different properties among the MHC I, IIA, and IIAX fibers. Future studies using electron microscopy should offer valuable insights into this possibility.

In contrast to contraction, during relaxation myosin ATPase is decoupled from actin and reflects its intrinsic ATPase domain function. The small differences in relaxed ATPase activity among fiber types would suggest that the ATPase domains are very similar, in sequence and/or in structure, among the different MHC isoforms. How the protein folds based on its sequence may impact its hydrolysis rate, phosphate release or ADP release rate, or how well the subunits interact with each other via electrostatic forces. Indeed, the small differences in ATPase rate among MHC isoforms are supported by the low structural variability among their myosin head region that contains the ATPase domain (1). The differences among SRX properties may be due to the high structural variability among the three loop regions of the MHC isoforms (1), which may play a role in the stabilization of the auto-inhibited state. Additionally, MHC I and IIA fibers having similar SRX ATPase rate agrees with their functional distinction from other MHC II isoforms, wherein MHC I and IIA fibers contract more slowly (3).

In conditions of constant passive stress, there is evidence of a strong positive correlation between a single fiber’s SRX fractional change and its sarcomere length change (5) that is beyond the maximal length examined in the present study. Here, we show the relaxation states’ fractional dependence on resting sarcomere length among multiple fibers. Longer resting sarcomere lengths resulted in more myosin heads being in the SRX state and decreased lifetimes of DRX and SRX (**Fig. 3*A-D***). As our sarcomere length range was only 1 μm, a small length difference appears to correlate with differing properties of relaxed myosin. Low spatial hindrance on the thick filament with increasing resting fiber sarcomere length would support our observation of greater SRX fraction (**Fig. 3*B***). This dependence may be due to decreased lattice spacing (6) or to altered interactions with stretch-sensitive myofilament proteins, such as myosin binding protein-C (24), as SRX has been shown to primarily occur in the C-zone of the thick filament (18). These altered interactions may induce activating changes in the ATPase domain in myosin that result in faster nucleotide turnover lifetimes in both relaxation substates (**Fig. 3*C*&*D***). Although our measurements cannot determine what is altering the properties of relaxed myosin with sarcomere length, they provide essential observations on the biochemical effects from small changes on fiber physical parameters.

Aside from preliminary measurements of SRX in five fibers from rabbit soleus muscle (27), SRX had not been previously studied in muscle considered slow-twitch and composed mainly of the MHC I isoform (29). Most previous skeletal muscle SRX measurements were conducted on rabbit psoas fibers (13, 27), which are predominantly MHC IIX (8), so the need to consider variable MHC expression was low. However, when rodent and human muscles are studied, fiber type information becomes important because of the diversity of MHC II subtypes (16, 25, 26).

In humans, MHC IIX fibers are extremely rare, expressing in less than 1% of the muscle fiber population (16). MHC IIX are more commonly coexpressed with MHC IIA, yielding “hybrid” MHC IIAX fibers, as seen in sedentary individuals (7, 10, 30). Therefore, determination of relaxation substates’ properties in human MHC IIX myosin depends on having a wide range of MHC IIX expression in the evaluated fibers. In the currently study, the range of MHC IIX expression was limited to 40-55%. There was no significant correlation between the fraction or lifetime parameters and the percent expression of MHC IIX in the fibers. Given that MHC IIX expression alters fiber cross-bridge kinetics (11, 14), there is interest in determining the relationship between relaxation substates and MHC IIX expression, which will be a focus of future studies.

In summary, we report that the myosin super-relaxed state in skeletal muscles is fiber-type dependent. Furthermore, by establishing the feasibility to quantify the population and lifetime of human skeletal muscle SRX, we set the stage for future biophysical studies examining the role of the relaxed states on human health, disease, and aging.

## Acknowledgement

We thank the University Imaging Center (http://uic.umn.edu) for technical support. This work was supported by grants from the National Institutes of Health to D.A.L. (R01AG031743), D.D.T. (R01AR032961, R37AG026160), L.A.P. (T32GM008244, T32AG029796, F30AG057108), M.S.M. (R01AG047245), and Dr. Jane Kent (R21AR073511) who generously supplied tissue for the young subject.

## List of Abbreviations

DRX: Disordered relaxed state
mantATP: 2’-/3’-O-(N’-Methylanthraniloy)-adenosine-5’-O-triphosphate
MHC: Myosin heavy chain
SRX: Super-relaxed state

